# A self-templated route to monodisperse complex droplets as artificial extremophile-mimic from coacervate-liposome interplay

**DOI:** 10.1101/2021.02.19.432011

**Authors:** Qingchuan Li, Qingchun Song, Wei Guo, Yang Cao, Youchuang Chao, Xinyu Cui, Jing Wei, Dairong Chen, Ho Cheung Shum

## Abstract

Bottom-up synthetic biology seeks to construct artificial cells from simple building blocks for exploring origin and principles of cellular life and material design. Although cellular life may have emerged spontaneously, programmable integration of building blocks into size, membrane property-controlled compartments (liposome or coacervate) towards cellular organization, without using specialized devices, has proven difficult. Here, we report a self-templated route to monodisperse complex droplets in bulk solution from coacervate-liposome synergy, with nanoliposome controlling coacervate size and coacervate templating on-surface nanoliposome fusion. Nanoliposome-coated monodisperse coacervates are self-assembled within 30 seconds, which are sealed by fusing nanoliposomes into size-controlled giant unilamellar vesicles (GUVs), with model building blocks combinatorially integrated into droplets. Defect-free membrane is established on coacervate, which render these complex GUVs surviving at extreme osmotic, salty and pH conditions (4 M NaCl, 100 mM HCl, 1 M NaOH), while providing homeostasis for enzymatic reactions, reminiscent of extremophiles.

## Main content

Cells manifest complexity in compositions and spatial organizations of compartments with a narrow size range. Bottom-up synthetic biology seeks to construct minimal living systems from biological or synthetic building blocks^1,2.^ Such efforts can help illuminate the origin and mechanism of cellular life^3,4^ and inspire cell-mimicking materials for advanced therapy^5-7^, microrobots^8,9,^ and biosensing^10^. Precision manipulation tools, such as microfluidics, have advanced this field towards the complexity exhibited by cellular life^11-17^. Facilitated by them, lipids and other building blocks can be sequentially assembled towards synthetic cells with controlled size, membrane lamellarity, and spatiotemporal organization of functional elements^18^. These have been challenging for bulk methodologies, although cellular life has originated, probably in the “primordial soup” of a complex organic mixture^19,20,^ without sophisticated tools. For example, lipid vesicle and the coacervate – a membraneless condensate droplet – have been two prevalent compartments for synthetic cells^3,21-23.^ Self-assembly of cell-sized monodisperse lipid vesicles or coacervates has rarely been achieved in bulk solution. Programmable organization of building blocks into droplets – like the combinatorial or sequential control afforded by microfluidics^18^ – has also been difficult using bulk methods. Bulk self-assembly’s lack of precision and programmability constitutes one of the important missing pieces of the puzzle as to how the inanimate has been integrated into primitive cellular life. Moreover, synthetic cells have recently attracted considerable interest for applications in therapy, biosynthesis, and biosensing^6,10,24,25^. Exploration of a self-assembly principle to bridge this gap in bulk solution will suggest the plausible route to cellular life from complex prebiotic mixtures and facilitate the emerging materials applications of synthetic cells by enabling mass production of customizable monodisperse droplets.

A recent finding shows the size of membraneless organelles (MLOs) in cells – condensate droplets similar to coacervates – can be controlled by adsorbed nanoscale protein clusters^26^; similar size control is also hypothesized from nano-sized vesicles, as indicated by a MLO at synapse coated by synaptic vesicles^27^. In addition, lipid vesicles are often utilized as precursors of a continuous membrane at an interface^18^, while similar membrane formation remains challenging on aqueous-aqueous interface, such as coacervate surface. In this work, we exploit nano-sized liposome as both size controller and membrane precursor for coacervate, and demonstrate a self-templated route to monodisperse microdroplets with controlled size, membrane lamellarity, and complexity in bulk solution. Briefly, liposomes serve to control the size of the generated monodisperse coacervates at surface, with building blocks combinatorially loaded into the droplets, and then fuse into a continuous membrane to form size-controlled giant unilamellar vesicles (GUVs) with controlled complexity. Two major challenges have been overcome to realize this self-templated route. First, different from oil-water and aqueous two-phase emulsion systems^28,29,^ the understanding of interfacial assembly on coacervates is still in its infancy^30^, and the utilization of particle adsorbers to generate monodisperse coacervates has been rarely achieved, either by liposomes or other particles^6,31-33.^ Hence, we perform a systematic study of coacervate formation in the presence of different particles or liposomes and establish the principles for monodisperse coacervate formation. Second, regarding to the challenge of membrane formation from nano-sized liposomes at aqueous-aqueous interface, we treat nanoliposome-coated coacervates through a freeze-thaw cycle – a possible physicochemical driver on early Earth^34^, with a defect-free membrane formed on coacervates. As an emergent consequence, the obtained defect-free monodisperse complex droplets can intriguingly survive and function at extreme pH, salt, or osmotic environments, a feat that has been impossible for either coacervate or liposome alone. Together, from the interplay of coacervate and liposome, we establish the programmability and precision for self-assembly in bulk solution towards monodisperse complex droplets with controlled integration of building blocks and robust functions.

### Assembly of monodisperse coacervates in nanoparticle solution

We first explore the possibility of monodisperse coacervate formation with liposomes or other nanoparticles as size controllers. Different from many earlier studies to assemble particles with as-prepared coacervates, we form coacervates in the presence of nanoparticles. This allows particles to affect coacervate formation at the onset of droplet nucleation. To form coacervates constituted by positively charged poly(diallyldimethylammonium chloride) (PDDA) and negatively charged adenosine 5-triphosphate (ATP), we first mix aqueous solutions of PDDA and particles, and subsequently ATP via vortexing for 30 s (Fig. 1a). In most of the particle-coacervate assembly scenarios, depending on PDDA and ATP molecular ratio *φ* and particle types, particles are extruded/recruited by the coacervate phase or adsorbed on the coacervate surface, with polydisperse coacervates formed (Cases 1-3, Fig. 1a-d and S1-3). By contrast, nanoparticle-stabilized monodisperse coacervates (Case 4) are demonstrated under specific conditions, such as 100 nm charged liposomes composed of neutral 1,2-dioleoyl-*sn*-glycero-3-phosphocholine (DOPC) and negatively charged 1,2-dioleoyl-*sn*-glycero-3-phospho-L-serine (sodium salt) (DOPS) with DOPS percentage *P* ranging from 5% to 12.5%. Other particles, including charged 100 nm liposomes of other compositions, silica particles encompassed by lipid membrane, magnetic nanoparticles, and some polystyrene (PS) particles, are all shown to co-assemble with PDDA and ATP to form monodisperse coacervates (Fig. 1b and S1). Moreover, except for PDDA-ATP, monodisperse coacervate formation is also observed for other coacervate constituents (Fig. 1c and S3), including PDDA-adenosine 5-diphosphate sodium salt (ADP), PDDA-poly(acrylic acid) (PAA), poly(allylamine hydrochloride) (PAH)-ATP, and poly(ethyleneimine) (PEI)-ATP. Together, these results demonstrate the universality of monodisperse coacervate formation in diverse coacervate-nanoparticle hybrid systems.

**Figure 1.**
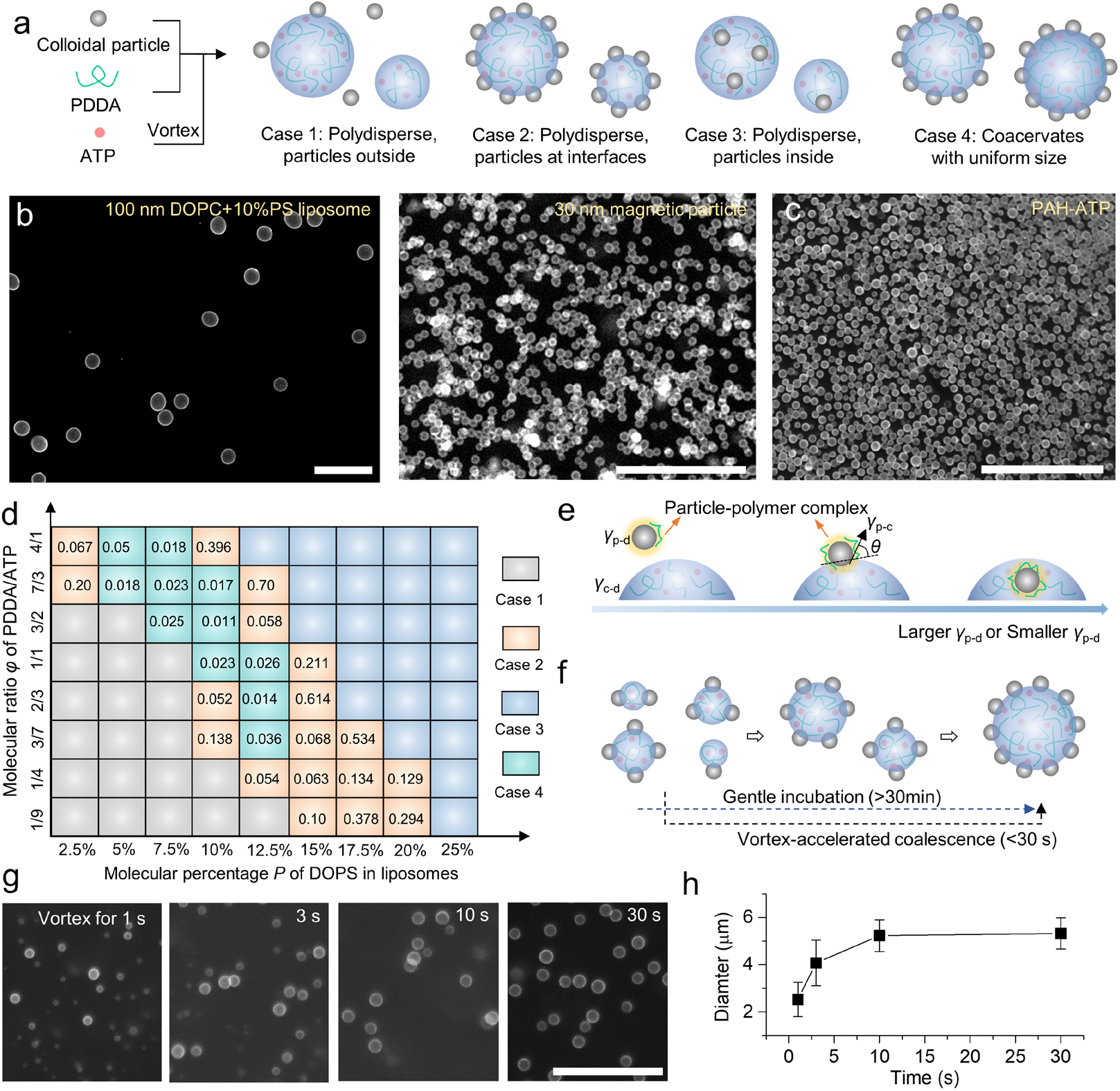
Vortex-promoted rapid assembly of monodisperse coacervates. **a**, Schematic showing the four cases after mixing PDDA, colloidal particles and subsequently ATP via vortex. **b**, Fluorescence images of monodisperse PDDA-ATP coacervates coated by 100 nm DOPC/DOPS (DOPS percentage *P* = 10%) liposomes or 30 nm magnetic nanoparticles. **c**, Fluorescence images of monodisperse PAH-ATP coacervates. **d**, Phase diagram for assembly in DOPC/DOPS liposome-PDDA-ATP system at different *P* and *φ*. Values in the table are calculated polydispersity index (PDI) from at least 100 coacervates for each case. Coacervate dispersion with PDI ≤ 0.05 is considered as monodisperse. **e**, Schematic illustrating particle partitioning across coacervate as a balanced particle-polymer complex. **f**, Schematic showing rapid formation of monodisperse coacervates from vortex-accelerated-limited-coalescence. **g**, Fluorescence images of PDDA-ATP (*φ*=3/2) coacervates coated by DOPC/DOPS (*P* = 10%) liposomes with vortexing time. h, Diameter of coacervates with vortexing time. The error bars represent the standard deviation. For all cases, *n*>100. The scale bars are 50 μm.

### Assembly of nanoparticles across coacervates

Coacervate-particle-interplay is ubiquitous in both biological and material sciences^26,31,35^. From our experiments where coacervates are assembled with diverse particles, we summarize several features of the coacervate-nanoparticle system. First, nanoparticles with higher charge density, such as liposomes containing more charged lipids (Fig. 1d) or particles with larger absolute particle zeta potential |*ζ*| (Fig. S4), prefer more coacervate phase. Second, the stoichiometry of coacervate constituents influences particle partitioning tendency. Negatively charged nanoparticles favor more coacervate phase with increasing ratio of cationic constituents to anionic constituents of coacervates. For instance, in PDDA-ATP-DOPC/DOPS liposome system, liposome favors coacervate phase with increasing PDDA/ATP ratio *φ*; similar results are also observed for PDDA/PAA, PDDA/ATP, PEI/ATP, and PAH/ATP coacervates (Fig. S3). Third, the electrostatic interaction strength between cationic and anionic coacervate constituents contributes to particle partitioning tendency. With fixed cationic component of coacervate, negatively charged nanoparticles tended to be recruited by coacervates with weaker electrostatic interaction strength between their constituents. Taking DOPC/DOPS liposomes with DOPS percentage *P*=10% as example, the liposomes assemble on the surface of PDDA-ATP (*φ*=3/2) coacervates (Fig. 1d) but are recruited by PDDA-ADP coacervate phase (PDDA more weakly associates with ADP than ATP) (Fig. S3).

These complex configurations of nanoparticle across coacervates can be rationalized by a particle-polymer complex (PPC) model. Due to adsorption of oppositely charged polymers, charged nanoparticle in coacervate system can be regarded to exist as a complex of particle and adsorbed polymers (PPC) (Fig. 1e). The electrostatic complexation between anionic and cationic molecules in coacervates has been reported to contribute to the interfacial tension between the coacervates and dilute phases *γ*_c-d_, which increases with electrostatic interaction strength^36^. Similarly, electrostatic complexation between polymers and nanoparticles is expected to contribute to the interfacial tensions of PPC with dilute phase (*γ*_p-d_); higher *γ*_p-d_ would be induced for more or stronger polymer adsorption on nanoparticles. Moreover, the difference in electrostatic interaction strength within PPC and coacervate leads to the interfacial tension *γ*_p-c_ between them, which can be anticipated to be *γ*_p-c_=*γ*_c-d_-*γ*_p-d_ when only tension from electrostatic complexation is considered.

Analogous to particle assembly in other emulsion systems^37^, the assembly of PPC across coacervate is then a result of the balance among *γ*_p-d_, *γ*_p-c_ and *γ*_c-d_, with PPC and coacervate interacting with contact angle *θ* following the Young’s equation^37^, cos(*θ*)=(*γ*_p-c_-*γ*_p-d_)/*γ*_c-d_=1-2*γ*_p-d_/*γ*_c-d_. The increase of *γ*_p-d_ or decrease of *γ*_c-d_ will facilitate larger *θ, i*.*e*., the partitioning preference of particle from dilute phase (*θ*=0°) to coacervate phase (*θ*=180°). Therefore, in PDDA-ATP-liposome system, with the increase of *P* or *φ*, which facilitates larger *γ*_p-d_ due to more or stronger adsorption of PDDA on liposome (Fig. S5), we observe partitioning tendency of liposomes from dilute phase to interface, and then coacervate phase (Fig. 1d). Moreover, the PPC model also explains the more preference of same charged liposomes for the coacervate phase of PDDA-ADP than PDDA-ATP (Fig. 1d and S3), in consideration of the smaller *γ*_c-d_ of PDDA-ADP coacervate from weaker electrostatic strength between coacervate constituents. Hence, the PPC model captures the diverse experimental observations in different coacervate-particle systems. More importantly, this model highlights the essential role of PPC in complex coacervation processes, which modulates structures of liposome-coacervate complexes and provides new paths towards programmable self-assembled structures.

To make PPC effective surface stabilizer for monodisperse coacervate formation, as indicated by particle assembly in other emulsion systems^37^, a sufficiently large contact angle *θ* ≤ 90° of PPC with coacervate should be established. This indicates *γ*_p-d_≤0.5*γ*_c-d_ according to the Young’s equation in coacervate-nanoparticle system. Consequently, a good balance between particle charge and coacervate composition is required, resulting in a narrow range of conditions for monodisperse coacervate formation (Fig. 1d). Liposome has shown to be a general stabilizer for generation of monodisperse coacervates of different components, as facilitated by the feasibility to modulate their charge states from lipid compositions to achieve the balance.

### Vortex-promoted rapid assembly of monodisperse coacervates

With the charged states of liposomes rendering them effective surface stabilizers, a limited-coalescence-process^28,29,^ as indicated by real-time observation of PDDA-ATP coacervate growth (Fig. S6), accounts for the size-controlled coacervate formation (Fig. 1f). Small coacervates with loosely packed liposomes on surface are firstly formed upon mixing of liposomes and charged molecules. These coacervates coalesce, consequently increasing liposome packing density on coacervate until further coalescence is inhibited. An incubation time over 30 min is generally required to complete the coalescence process, which results in coacervates with compromised monodispersity (Fig. S6). However, vortexing is found to significantly accelerate coalescence process and promote droplet monodispersity. As a result, monodisperse coacervates (Polydispersity index≈0.01) are obtained in bulk solution within only 30 s (Fig. 1g, h), with tunable diameter from several micrometers to tens of micrometers by adjusting the amount of particles used (Fig. S7). Vortexing can promote the collision of immature coacervates, which accelerates coalescence and avoids insuffient coalescence of immature coacervates by overcoming electrostatic repulsion among coacervates, consequently resulting in better monodispersity of coacervates.

### Spatiotemporal control of the complexity of monodisperse coacervates

The observed limited-coalescence process and sensitivity of coacervate coalescence to vortex during monodisperse coacervate formation (Fig. 1g) inspire a combinatorial approach to engineer the complexity of coacervate droplets. To this end, a series of small precursor coacervates (1)-(9) with various structures are first formed via gentle mixing of PDDA, ATP, liposomes with red (R-liposome) or green (G-liposome) fluorescence, PAH, or PS particles with blue (B-PS particle) or green (G-PS particle) fluorescence (Fig. 2a). These precursor coacervates are then combinatorically mixed and vortexed for their fusion, yielding hierarchical coacervates with elaborately engineered structures (Fig. 2b-j). On coacervate surface, R-liposomes and G-liposomes can be controlled to form a homogeneous mixture or jammed, heterogeneous rafts (Fig. 2b, c and S8). Inside coacervates, we engineer the spatial organization of G-liposome and B-PS particle with various configurations (Fig. 2d-g). Multiphase coacervates comprising PDDA-ATP rich phase and PAH-ATP rich phase (Fig. 2h and S9) are generated via self-fusion of coacervate building blocks (7); the resultant morphology resembles those of eukaryotic cells, with inner and outer phases mimicking the nucleus and cytoplasm respectively. An even higher degree of complexity can be introduced into these multiphase droplets by loading B-PS particle or G-PS particle as “organelles” (Fig. 2i-j). This combinatorial method is demonstrated to effectively organize matter into diverse configurations of monodisperse coacervates.

**Figure 2.**
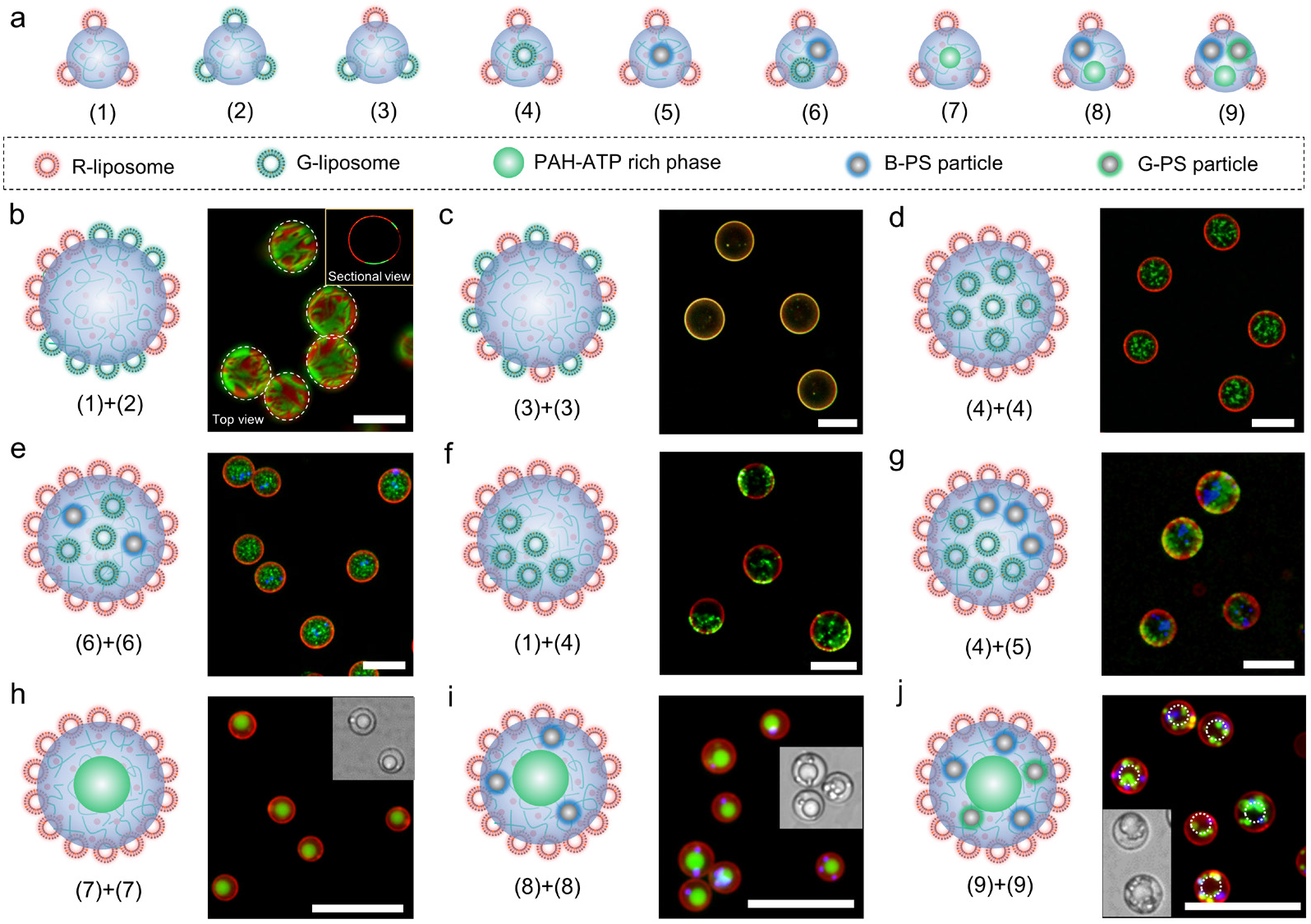
Assembly of monodisperse coacervates with defined complexity via combinatorial fusion. **a**, Schematic of the nine small coacervate building blocks (1)-(9) formed via the gentle mixing of PDDA, ATP, PAH or different particles for combinatorial fusion experiments. R-liposome and G-liposome indicate liposomes with respectively red and green fluorescence. B-PS particle and G-PS particle indicate PS particles with blue and green fluorescence respectively. **b-j**, Schematic and fluorescence images of various hierarchical coacervates formed via vortex-promoted fusion of selected coacervate building blocks from **a**. The scale bars are 20 μm.

### Bulk assembly of size-controlled GUVs

Besides serving as surface stabilizer to control the size and complexity of coacervates, liposomes can further be precursors of a continuous lipid membrane on coacervates through a freeze-thaw procedure, leading to monodisperse complex GUVs. We freeze liposome-coated monodisperse coacervates (L-coacervates) in liquid nitrogen or -80°C/-20°C refrigerators, and then allow them to thaw at room temperature. Coacervate-supported GUVs (CGUVs) are generated from liposome rupture by freezing and subsequent fusion of membrane patches during thaw (Fig. 3a). In particular, monodisperse CGUVs (PDI<0.02) are obtained by freezing L-coacervates in liquid nitrogen or -80°C, achieving a transformation efficiency of > 50% from L-coacervates (Fig. S10). The membrane on CGUVs displays tunable permeability to small organic molecules such as fluorescein. With PDDA in dispersion media, fluorescein can permeate through membrane and be enriched in the coacervate core. After washing the CGUVs with excess 5 mM ATP plus 100 mM NaCl solution to remove the adsorbed PDDA, membrane permeability to fluorescein is blocked (Fig. 3b). We incubate the non-leaky CGUVs with melittin and then fluorescein. The coacervate phase of all CGUVs containing melittin emits fluorescence of fluorescein, indicating membrane unilamellarity^14^. The phospholipid membrane and coacervate phase in CGUVs are in fluidic phase, as evidenced by rapid fluorescence recovery of fluorescein in fluorescence recovery after photobleaching (FRAP) experiments (Fig. 3c, d).

**Figure 3.**
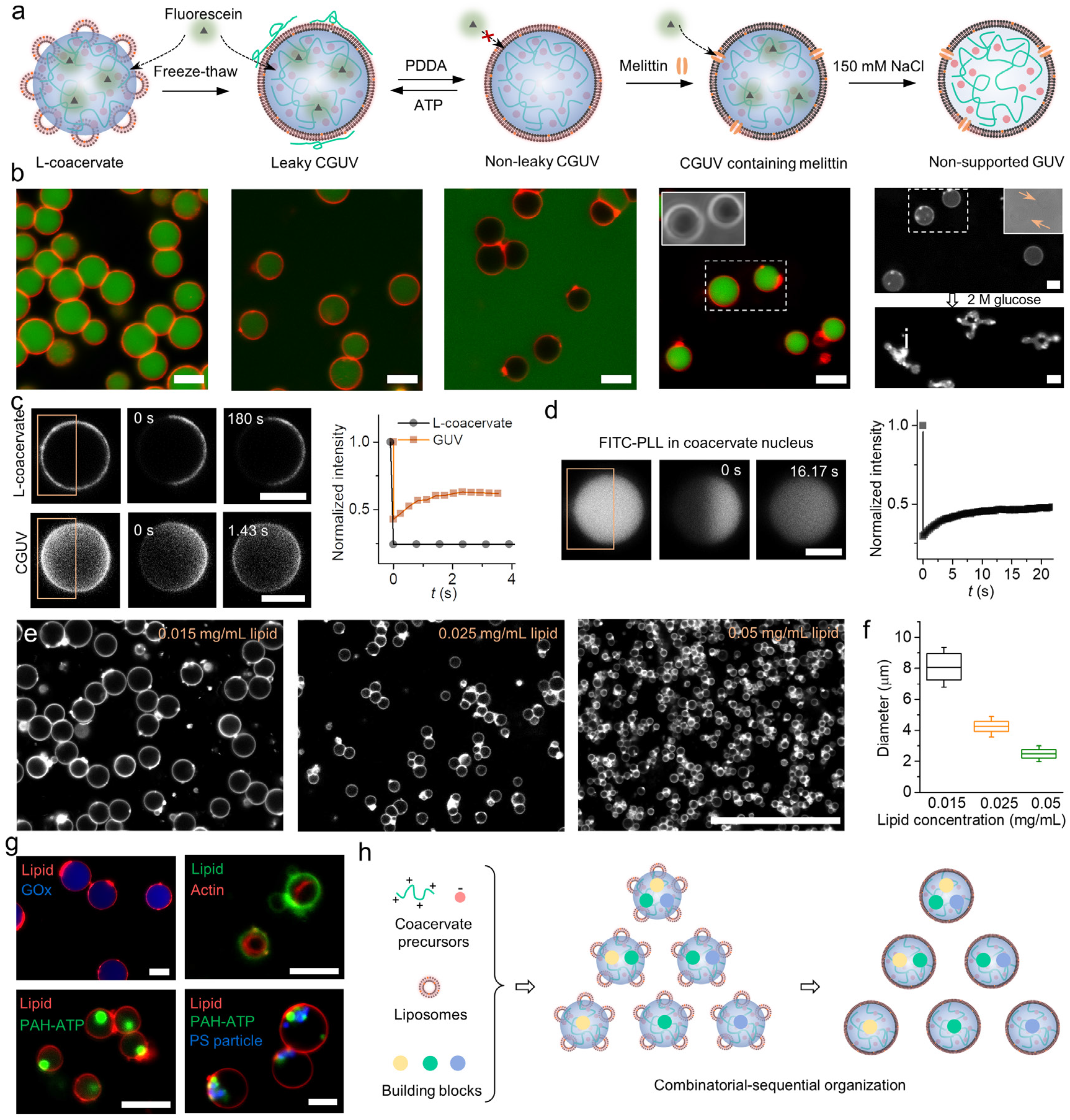
Bulk assembly of size-controlled GUVs. **a**, Schematic illustrating the formation and permeability test of DOPC/DOPS/cholesterol (37/3/10) CGUVs on PDDA-ATP coacervates (*φ*=7/3) (CGUVs) from liposome-coated coacervates (L-coacervate) and the subsequent disassociation of coacervate core to generate non-supported GUVs. **b**, Fluorescence images showing the permeability to fluorescein of L-coacervate, leaky CGUV, non-leaky CGUV, and CGUV containing melittin (from left to right) and the deformation of non-supported GUVs under osmotic stress. The insets are phase contrast images of droplets in the yellow dashed boxes indicating the existence or disassociation of coacervate core. **c** and **d**, Fluorescence recovery after photobleaching (FRAP) experiments of rhodamine-DOPE in L-coacervate and CGUV (**c**) and fluorescein isothiocyanate-labelled poly-L-lysine (FITC-PLL) inside CGUV (**d**). **e** and **f**, Size control of CGUVs formed via quick freezing in liquid nitrogen. The error bars represent the standard deviation. For all cases, *n*>100. **g**, Fluorescence images of complex CGUVs encapsulated with GOx, actin, PAH-ATP rich phase, or B-PS particles. h, Schematic illustrating the programmable organization of a starting mixture containing coacervate precursors, liposomes, and other building blocks from coacervate-liposome interplay. Scale bars in **b-d, g** are 5 μm. Scale bar in **e** is 50 μm.

The CGUVs retain the size and complexity of their mother L-coacervates. CGUVs with average diameter from ∼8 μm to ∼2 μm can be generated when lipid concentration in mixtures varies from 0.015 mg/mL to 0.05 mg/mL (Fig. 3e, f). By tailoring complexity of mother coacervates with the combinatorial strategies (Fig. 2), down-stream complex CGUVs encapsulating glucose oxidase (GOx), actin, PAH-ATP rich phase, or B-PS particles are obtained (Fig. 3g). Moreover, following assembly, the coacervate nucleus can be disassembled by incubating CGUVs in 150 mM NaCl solution containing melittin, resulting in non-supported GUVs (Fig. S11). These GUVs exhibit active membrane fluctuations and can deform into lipid tubes in hyperosmotic glucose solution (Fig. 3b), analogous to GUVs formed by traditional electroformation methods^38^.

The coacervate surface has been previously employed as reaction site for lipid self-assembly^3,39,40^. Our CGUVs are unique in following aspects: First, self-assembly of monodisperse GUV with an aqueous core is demonstrated for the first time in bulk solution. Second, different from previous studies^3,24^,25, a defect-free lipid bilayer is established on coacervates, as evidenced by the non-permeability to fluorescein (Fig. 3b). Third, our approach can be potentially applied to assemble a continuous, defect-free membrane from cell-derived vesicles on coacervates. This has been challenging for current membranization strategies^3,39^ but may open new possibilities for biomedical or biomimetic applications^41^. Finally, and most importantly, a self-templated route to membrane-bound droplets with controlled size, membrane lamellarity and complexity is established by combining the combinatorial self-assembly of building blocks into monodisperse coacervates and sequential membrane formation (Fig, 3h). Hence, from the interplay between coacervates and liposomes – the two coexisting compartmentalization entities for cells or probably primitive life, we establish the programmability and precision for bottom-up synthetic biology in bulk solution.

### Synthetic “extremophiles” survive and function at extreme environments

A structural and functional stability at extreme environments is observed on the monodisperse CGUVs with defect-free membrane from our self-templated route. GUVs have been well-known to be sensitive to small osmotic pressure to deform^38^. Moreover, coacervates are generally sensitive to salt or pH to disassemble, due to weakening of the electrostatic interactions that hold molecules together. However, astonishing stability to extreme osmosis, salt concentration or pH condition is observed for CGUV, the hybrid structure of defect-free GUV and PDDA-ATP coacervate core (Fig. 4a). The CGUVs retain spherical morphology in 100 mM MgCl_2_, 1 M glucose, 1.5 M NaCl, 100 mM HCl, and 1 M NaOH solution. According to turbidity tests, PDDA-ATP coacervate disassemble with NaCl concentration above 50 mM, while CGUVs are stable even when NaCl concentration reaches 4.0 M, which indicates a high osmotic pressure of ∼100 atm. Regarding pH sensitivity, PDDA-ATP coacervates disassemble at pH<3 or NaOH concentration > 100 mM, while CGUVs remain stable at acidic condition at pH<2 and alkaline condition with NaOH concentration > 1 M. Hence, an ultrastable droplet structure is demonstrated by integrating two well-known fragile structures.

**Figure 4.**
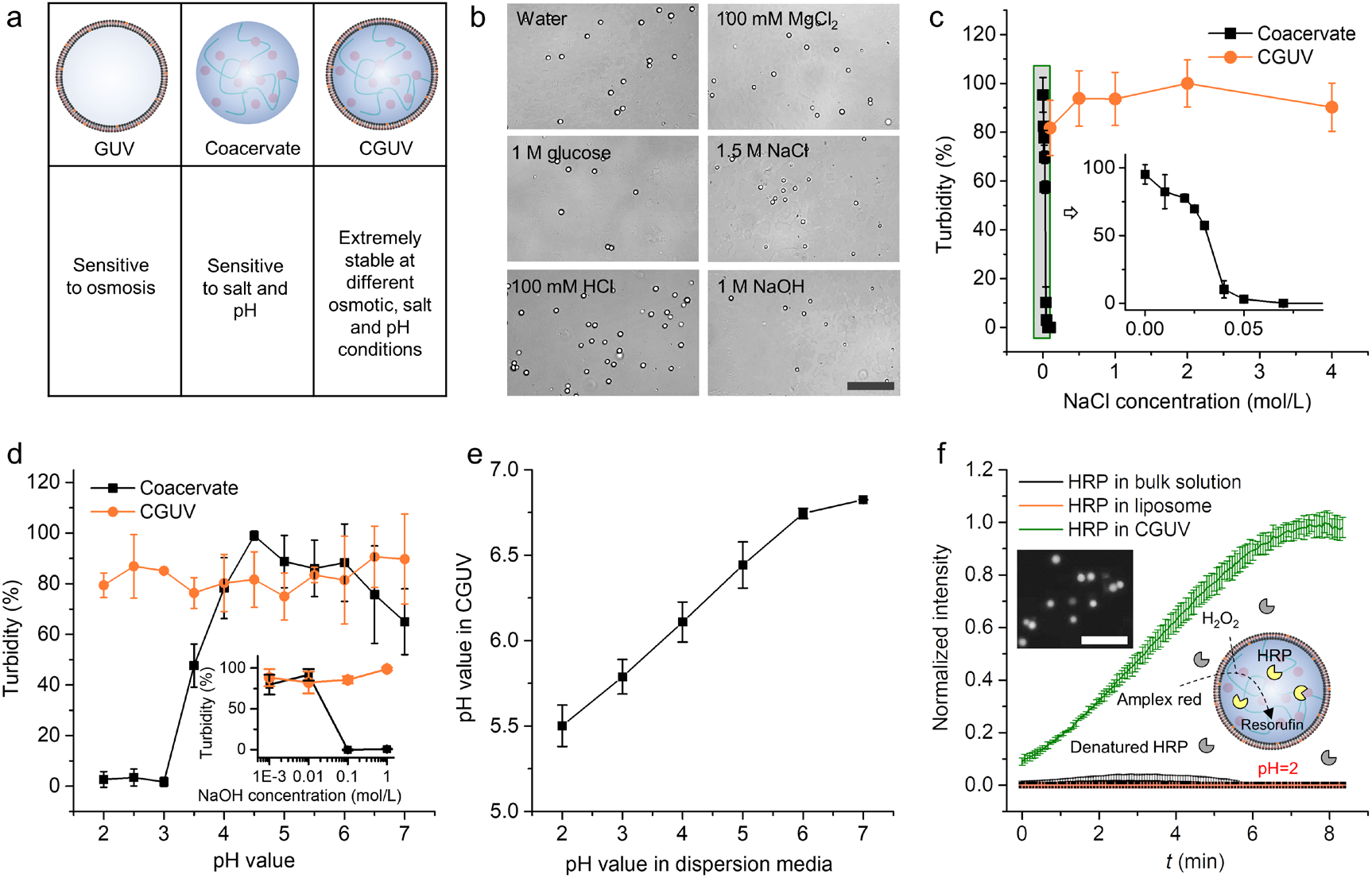
Stability and function of CGUVs at extreme environments. **a**, Schematic showing that CGUV exhibits extreme stability, which cannot be achieved either by GUV or coacervate. **b**, Bright field images CGUVs are stable at different solution environments. **c**, Stability of coacervate and CGUV with NaCl concentration from turbidity test. **d**, Stability of coacervate and CGUV at different pH conditions. **e**, The apparent pH value in CGUV with pH value of the dispersion media. **f**, The CGUV keeps the activity of HRP even in an acidic dispersion solution that denatures HRP in bulk solution and liposome. The inset shows a schematic of enzymatic reaction and a fluorescence image of fluorescent resorufin products in CGUVs. The scale bars are 20 μm.

The stability of CGUV is an outcome of the synergy between the defect-free membrane and coacervate core. The defect-free phospholipid membrane blocks the entry of ions to disassemble coacervate, leading to stability of CGUVs at high salt concentrations. In turn, the viscous coacervate core imparts CGUVs mechanical stability to balance high osmotic stress. Moreover, the CGUV contains high concentration of ATP, which is anticipated to buffer the microdroplet and induce the pH stability (Fig. 4d). We estimate the pH value within CGUV by using pH-sensitive FITC-PLL as probe, the logarithmic fluorescence intensity of which has been shown to linearly reduce with pH decrease (Fig. S12). As shown in Fig. 4e, the pH value in CGUV is between 5.5 to 7 when pH value outside varies from 2-7, indicating pH homeostasis^42^ within CGUV to render them the pH stability (Fig. 4d). Moreover, the pH homeostasis enables the CGUVs enzymatic microreactors at extreme pH environment. In a solution media of pH=2, horseradish peroxidase (HRP) in bulk solution or liposomes denatures. However, HRP in CGUVs retains high catalytic ability to transform amplex red into red fluorescent resorufin (Fig. 4f). These demonstrate the robustness of these monodisperse, defect-free CGUVs from our self-templated route to adapt and execute functions at extreme conditions, reminiscent of extremophiles at harsh environments^43^.

## Conclusions

This work demonstrates a self-templated route to monodisperse droplets with controlled size, membrane lamellarity, and programmable complexity in bulk solution from coacervate-liposome interplay. Nano-sized liposome is shown to be a general size controller for coacervates on surface; in turn, coacervates serve as the substrate for fusion of nano-sized liposomes into a continuous membrane through a freeze-thaw procedure. Consequently, self-assembly of monodisperse coacervates, GUVs, and CGUVs in bulk solution are demonstrated for the first time, with different building blocks combinatorially organized into droplets. Furthermore, the coacervate-liposome interplay imparts the monodisperse, defect-free CGUVs structural and functional stability under extreme osmotic, salt, and pH conditions that is otherwise unachievable for either coacervates or liposomes alone, suggesting the concept of a synthetic “extremophile” model working at extreme environments.

The self-templated route establishes the precision and programmability towards cell organization in bulk solution. It may have played a role in integrating prebiotic building blocks towards cellular life. Different from previous focus on exploring the integrated complexity to be achieved within individual droplets^3,6,18,^ the self-templated route allows for programmable organization in a complex molecular mixture. Building blocks can be combinatorially organized into diverse protocells, from some of which the integration of compartmentalization, replication, and metabolism essential for cellular life may have achieved. Moreover, the prebiotic environments on early Earth could be harsh, and the CGUVs may have provided a plausible microenvironment with superior stability for primitive metabolic processes. Hence, the self-templated route provides a new view about how coacervates and lipid vesicles, when co-existed in a “primordial soup”, may contribute synergistically to origin of life.

The self-templated route enables scalable customization of monodisperse coacervates, CGUVs and non-supported GUVs with controlled properties. Unlike microfluidic methods^14^, this approach does not require delicate control over manipulating liquids. Monodisperse coacervates or GUVs can be generated quickly in bulk solution with high yields. The ease for mass-production, with precise control of droplet size and composition, will facilitate the emerging application of synthetic cell products for therapeutic, biosensing or biosynthesis^6,10^,25. Moreover, the stability observed on the defect-free CGUVs at extreme environments is promising to extend the application scope of synthetic cells towards conditions that would normally destabilize liposomes, coacervates, or enzymes, for example, in the acidic environment of stomach for therapy or in wastewater for detection or degradation of pollutants.

## Methods

### Monodisperse coacervate formation in colloidal solution

Complex coacervates were assembled from phase separation of counter-charged molecules in the presence of nanoparticles. In PDDA-ATP-nanoparticle system, PDDA stock solution (50 mM), particle dispersion, and water were mixed firstly, and then subsequently with ATP stock solution (50 mM) via vortexing. The final volume of the mixture was 100 μL, with the concentration of PDDA, ATP, and nanoparticles modulated via controlling the volume of stock solution used. Different nanoparticles, including liposomes of different compositions, PS particles, silica particles, magnetic nanoparticles, and proteins were used to co-assemble with coacervates. The assembly results, including the partitioning tendency of particles across coacervates and monodisperse coacervate (polydispersity index PDI<0.05) formation, were characterized using fluorescence microscope. The influences of PDDA or ATP concentration, the molecular ratio *φ* of PDDA and ATP, and particle types on assembly results were investigated. Moreover, the effect of coacervate constituents, including PDDA/ADP, PDDA/PAA, PAH/ATP, and PEI/ATP, on assembly results was also studied by using DOPC/DOPS liposomes as model particles. The principles guiding the partitioning tendency of particles across coacervates and monodisperse coacervate formation were summarized from these assembly experiments with different nanoparticles and coacervate composition.

### Combinatorial control of the complexity of monodisperse coacervates

A combinatorial approach was developed to engineer the complexity of coacervates. A series of small coacervate precursors was firstly prepared by gently mixing PDDA stock solution, particles (DOPC/DOPS liposomes with respective red and green fluorescence, 0.5 μm PS-COOH particles with green fluorescence, or 1 μm PS-COOH particles with blue fluorescence), or PAH stock solution, and subsequently with ATP stock solution. Selective coacervate precursors were then mixed and vortexed for 30 s to promote their coalescence, resulting in monodisperse coacervates with controlled complexity.

### Assembly of monodisperse GUVs

We used a freeze-thaw method to generate size-controlled GUVs from liposome-coated coacervates (L-coacervate). The sample of monodisperse PDDA-ATP Coacervates (*φ*=7/3) coated by 100 nm DOPC/DOPS/Chol (molecular ratio=37/3/10) liposomes was frozen at -20°C, -80°C, or in liquid nitrogen and then allowed to thaw at room temperature. NaCl was then added (Final concentration>150 mM) to disassemble coacervates that were not encompassed by a continuous membrane. The turbidity difference before and after adding NaCl was used to calculate the transformation efficiency from L-coacervates to CGUVs. The morphology of obtained CGUVs was characterized under the fluorescence microscope. The influence of freezing conditions (in liquid nitrogen, or at -80°C or -20°C) on transformation efficiency and CGUV morphology was investigated. Membrane permeability to small molecules was investigated by incubating the CGUVs with NaCl or fluorescein. Membrane unilamellarity was verified by checking membrane permeability to fluorescein after incubating CGUVs with 5 μg/mL melittin. Fluidity of membrane and coacervate phase was characterized through fluorescence recovery after photobleaching (FRAP) experiments. Non-supported GUVs without the coacervate core were further obtained by incubating CGUVs in solution containing 5 μg/mL melittin and 150 mM NaCl. The size and complexity of GUVs were modulated by varying those of mother L-coacervates.

### CGUVs at extreme environments

The structural stability of CGUVs at extreme osmotic, salt, and pH conditions was checked from microscope images and turbidity test of CGUVs at different NaCl, MgCl_2_, glucose, NaOH, and HCl concentrations. To evaluate the ability of CGUVs to function at extreme environment, HRP (0.5 U/mL) was co-assembled with PDDA, ATP, and liposomes, and frozen and thawed to form CGUVs encapsulated with HRP. These CGUVs containing HRP were then dispersed in solution media with pH value = 2 that would generally denature HRP and incubated for 60 min. The catalytic ability of HRP in CGUVs was checked by the quantaRed™ enhanced chemifluorescent HRP substrate kit using multi-mode microplate reader. As control, the catalytic ability of HRP (0.5 U/mL) in bulk solution and liposomes with pH=2 of dispersion media was also checked.

## Data availability

All data that support the findings of this study are available within the article and its Supplementary Information, and/or from the corresponding author upon reasonable requests.

## Acknowledgments

This research is supported by the General Research Fund (Nos. 17304017, 17305518, and 17306820) and National Natural Science Foundation of China/RGC Joint Research Scheme (N_HKU718/19), the NSFC Excellent Young Scientists Fund (Hong Kong and Macau) (21922816), as well as the Seed Funding for Strategic Interdisciplinary Research Scheme 2019/20 from the University of Hong Kong. Q.C.L. was supported by the Hong Kong Scholars Program. H.C. S. was partially supported by the Croucher Foundation through the Croucher Senior Research Fellowship.

## Author contributions

H. C. S., D. R. C., and Q. C. L. supervised the research. H. C. S., D. R. C., Q. C. L., and Q. C. S conceived and designed the experiments. Q.C.L., Q. C. S., J. W, Y. C., and X. Y. C. performed experiments. Q.C.L., Q. C. S., J. W, Y. C., Y. C. C., and X. Y. C. analyzed the data. H.C.S. and Q.C.L. wrote the paper, and all authors commented on the paper. Q. C. L and Q. C. S contributed equally to this work.

## Additional information

Supplementary information is available in the online version of the paper. Correspondence and requests for materials should be addressed to H. C. S., D. R. C., and Q. C. L.

