## Supplemental text and figures for "A self-templated route to monodisperse complex droplets as artificial extremophile-mimic from coacervate-liposome interplay"

**Experimental Section**

**Materials**

1,2-dioleoyl-*sn*-glycero-3-phosphocholine (DOPC), 1,2-dioleoyl-*sn*-glycero-3-phospho-L-serine (sodium salt) (DOPS), 1,2-dioleoyl-*sn*-glycero-3-phosphate (sodium salt) (DOPA), 1,2-dipalmitoyl-*sn*-glycero-3-phosphoethanolamine-N-(cap biotinyl) (sodium salt) (Biotin-lipid), 1,2-dioleoyl-3-trimethylammonium-propane (chloride salt) (DOTAP), 1,2-dipalmitoyl-*sn*-glycero-3-phosphoethanolamine-N-(7-nitro-2-1,3-benzoxadiazol-4-yl) (ammonium salt) (NBD PE), and 1,2-dioleoyl-*sn*-glycero-3-phosphoethanolamine-N-(lissamine rhodamine B sulfonyl) (ammonium salt) (Rhodamine PE) were purchased from Avanti Polar Lipids (USA). Poly(diallyldimethylammonium chloride) solution (PDDA, average $M_w$<100 kDa, 35% wt.% in water), Poly(ethyleneimine) solution (PEI, $M_w$~750 kDa, 50% (w/v) in water), Poly(allylamine hydrochloride) (PAH, $M_w$~17500), Poly(acrylic acid) (PAA, $M_w$~1800), Adenosine 5′-triphosphate disodium salt hydrate (ATP), Adenosine 5′-diphosphate sodium salt (ADP), Fluorescein isothiocyanate (FITC) labelled Poly-L-lysine (FITC-PLL), Poly(FITC allylamine hydrochloride) (FITC-PAH), iron oxide(II,III) magnetic nanoparticles solution (30 nm, carboxylic acid functionalized), amine-modified polystyrene (PS) nanoparticle (100 nm, fluorescent orange), cholesterol, ethanol, hydrochloric acid, sodium hydroxide, sodium chloride, glucose, sodium oleate, and fluorescein were purchased from Sigma. Horseradish peroxidase (HRP) was bought from Solarbio. QuantaRed™ enhanced chemifluorescent HRP substrate kit was purchased from ThermoFisher Scientific. Streptavidin coated PS nanoparticle (100 nm), 50 nm PS-COOH nanoparticle (fluorescent dragon green), 0.5 μm PS-COOH nanoparticle (fluorescent dragon green), and 1 μm PS-COOH nanoparticle (fluorescent glacial blue) bought from Bangs Laboratories, Inc. 100 nm silica particle was from Nanjing Nanorainbow Biotechnology Co., Ltd. Ag nanowire (30 nm diameter) was from Nanjing XFNANO Materials Tech Co., Ltd. 75 nm diameter PS-COOH nanoparticle (fluorescent green) was from BaseLine ChromTech Research Centre (China). Millipore Milli-Q water with a resistivity of 18.2 MΩ cm was used in the experiments.

The following stock solutions were prepared in water for coacervates formation: PDDA (50 mM, pH 7), PEI (50 mM, pH 7), ATP (50 mM, pH 7), ADP (50 mM, pH 7), PAA (50 mM, pH 7), PAH (50 mM, pH ~3). Liposomes of various compositions were formed through an extrusion method[1]. 1 mg phospholipid mixture was dried in a centrifuge tube. 1 mL of water was then added and vortexed. The mixture was extruded through polycarbonate filter with 100 nm pore size back and forth for 11 times. The composition of the liposomes was controlled by varying the ratio of different components in the phospholipid mixture. 0.2 wt% Rhodamine PE or NBD PE were incorporated for visualization under fluorescence microscope. Lipid bilayer coated silica particles were prepared via the incubation of liposomes (1 mg lipid) with 2 mg 100 nm silica particles in ultrasonic bath for 1h and subsequent removement of free liposomes via centrifugation. 30 nm magnetic particles, 1 μm magnetic particles and 30 nm diameter Ag nanowires were used as obtained. Other particles were diluted to 1mg/mL before use.

**Assembly of liposomes with coacervates with different constituents**

The assembly of particles with another four cationic specie-anionic specie pairs, namely, PDDA with two anionic species, ADP and PAA, and ATP with two cationic species, PAH and PEI, were investigated. DOPC/DOPS liposomes were taken as particle model. The stock solution of cationic specie, liposomes, water, and stock solution of anionic specie were mixed via vortex. The volume of the final mixture was 100 μL. For PDDA-ADP pair, the influence of PDDA/ADP molecular ratio $\varphi$(PDDA/ADP) and DOPS percentage $P$ in DOPC+DOPS liposomes were investigated. The sum concentration of PDDA and ADP in the final mixture was fixed at 5 mM. Liposome concentration was 0.10 mg/mL. For the assembly of PDDA-PAA pair with DOPC/DOPS ($P$=50%) liposomes, the influences of NaCl concentration and PDDA/PAA molecular ratio $\varphi$(PDDA/PAA) were investigated. The sum concentration of PDDA and PAA was 20 mM. Liposome concentration was 0.10 mg/mL. For the assembly of PAH-ATP pair with DOPC/DOPS ($P$=10%) liposomes, the influences of pH value in the final mixture and PAH/ATP molecular ratio $\varphi$(PAH/ATP) were investigated. [PAH]+[ATP] concentration was 25 mM. Liposome concentration was 0.10 mg/mL. For the assembly of PEI-ATP pair with DOPC/DOPS ($P$=10%) liposomes, the influence of pH value in the final mixture and PEI/ATP ratio $\varphi$(PEI/ATP) were investigated. [PEI]+[ATP] concentration was 10 mM. Liposome concentration was 0.05 mg/mL. The assembly results were characterized using fluorescence microscope and organized as phase diagrams.

**Characterization**

The fluorescence and bright field images were taken by Leica DMIL, LED, Fluo fluorescence microscopy, Zeiss Axio Observer A1 fluorescence microscope and Zeiss LSM710 laser confocal microscope. Zeta-potential of liposomes, particles, and liposome coated PDDA-ATP coacervates were measured using Zetasizer Nano Z (Malvern Instruments, UK). The pH value was measured using the PHSJ-3F pH meter from Shanghai INESA Scientific Instrument Co., Ltd. The turbidity and fluorescence intensity were measured in 96-well plates by SpectraMax iD3 Multi-Mode Microplate Reader (Molecular Devices, CA). The turbidity was calculated by measuring the absorbance at $\lambda$=500 nm. The fluorescence intensity of fluorescein at 516 nm was collected at excitation wavelength of 470 nm. The fluorescence intensity of resorufin at 590 nm was collected at excitation wavelength of 550 nm.

### Supplementary Figures

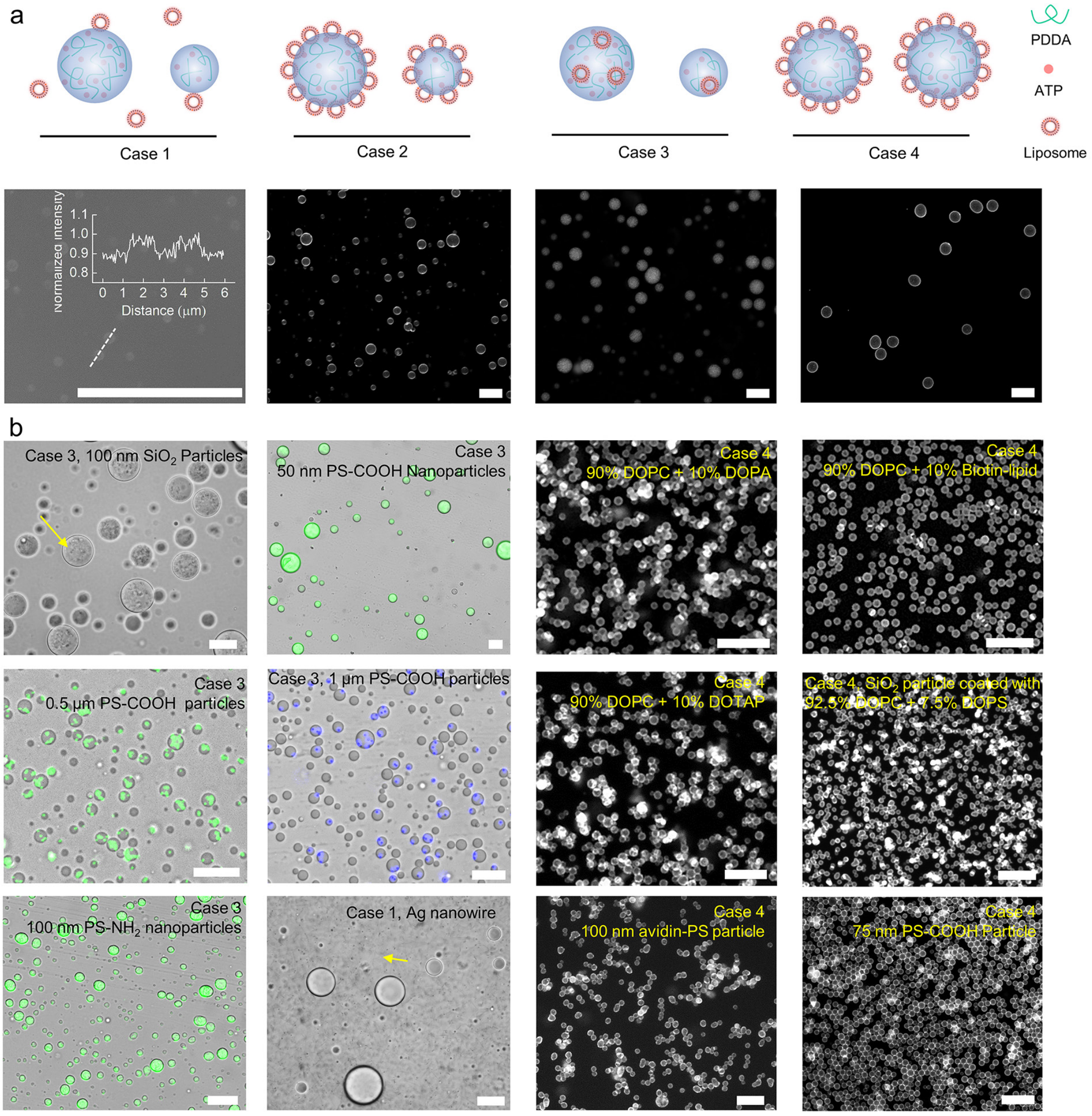


**Figure S1. Assembly of coacervates in colloidal solution. a,** Schematic and fluorescence images for the four assembly cases in DOPC/DOPS liposomes-PDDA/ATP coacervate hybrid system. The inset is the fluorescence profile along the white dash line. **b,** assembly configurations of other particles with PDDA/ATP coacervates. The yellow arrows indicate particles. The scale bars are 25 μm.

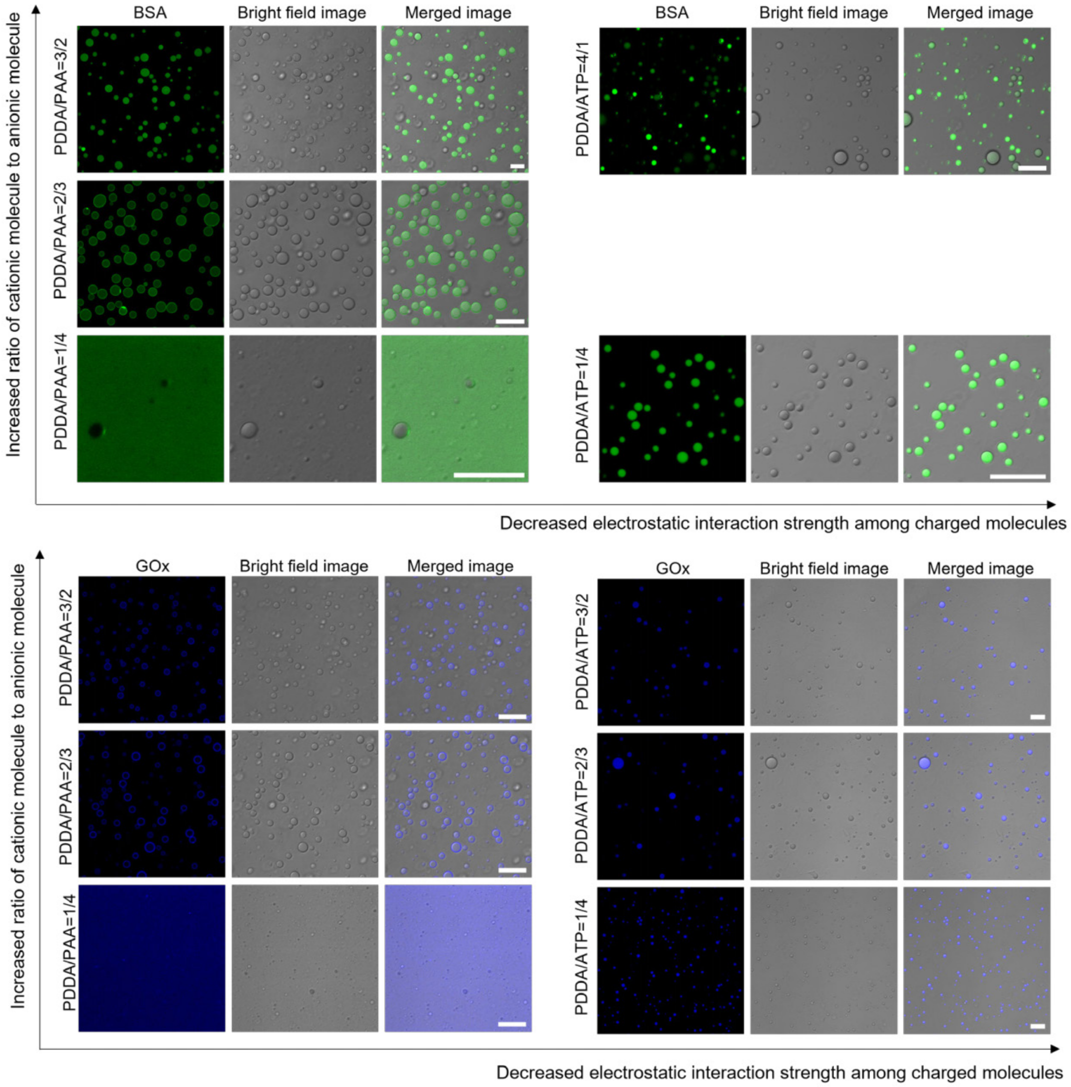


**Figure S2. Assembly of protein particles with coacervates.** In PDDA-ATP system, [PDDA]+[ATP]=5 mM. In PDDA-PAA system, [PDDA]+[PAA]=25 mM. 10% PEG 8000 is used as crowding reagent. The scale bars are 10 μm. PDDA and PAA are more strongly associated than PDDA and ATP. BSA and GOx located inside PDDA-ATP coacervates regardless of PDDA/ATP ratio. In PDDA-PAA system, the partitioning of BSA and GOx across coacervates depends on PDDA/PAA ratio. With the increase of PDDA/PAA ratio, protein particles exhibit a partitioning tendency from the dilute phase to coacervate phase.

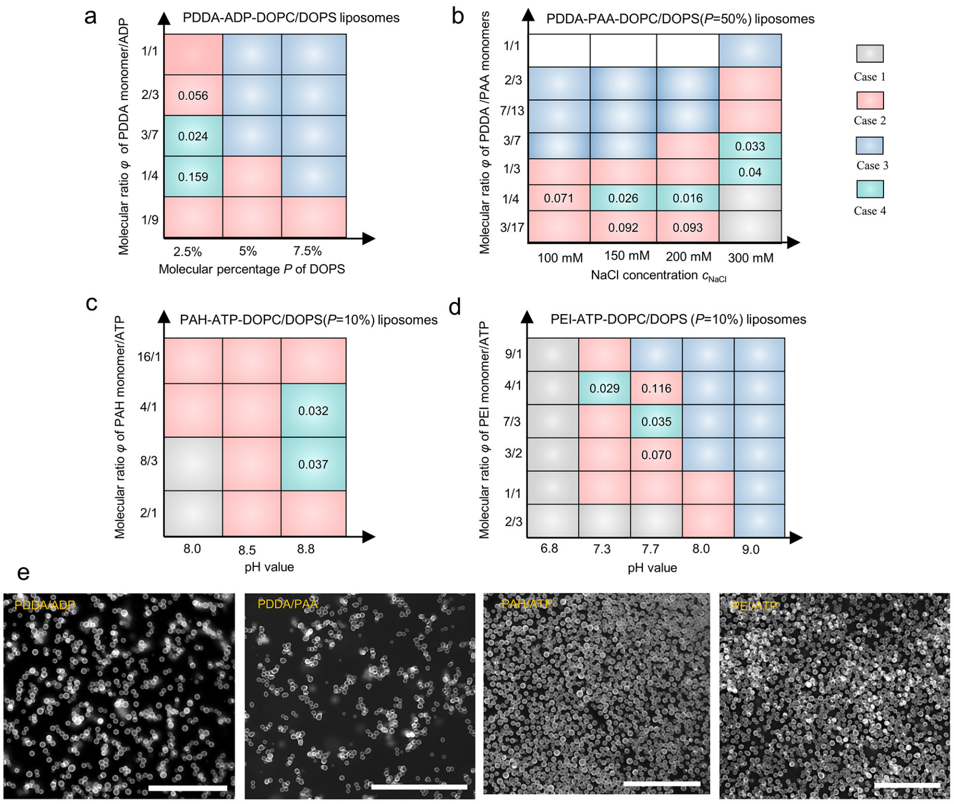


**Figure S3. Formation of monodisperse coacervates from other charged coacervate constituents. a**, Phase diagram showing the influence of *P* and [PDDA]/[ADP] ratio on coacervate formation. [PDDA]+[ADP] is 5 mM. Liposome concentration is 0.10 mg/mL. **b**, Phase diagram showing the influence of NaCl concentration and [PDDA]/[PAA] ratio on coacervates formation with the existence of DOPC/DOPS (*P*=50%) liposomes. [PDDA]+[PAA] is 20 mM. Liposome concentration is 0.10 mg/mL. **c**, Phase diagram showing the influence of pH value and [PAH]/[ATP] ratio on coacervate formation with existence of DOPC/DOPS (*P*=10%) liposomes. [PAH]+[ATP] is 25 mM. Liposome concentration is 0.10 mg/mL. **d**, Phase diagram showing the influence of pH value and [PEI]/[ATP] ratio on coacervate formation with the existence of DOPC/DOPS (*P*=10%) liposomes. [PEI]+[ATP] is 10 mM. Liposome concentration is 0.05 mg/mL. **e**, Fluorescence images of the monodisperse coacervates with different components. The scale bars are 50 μm.

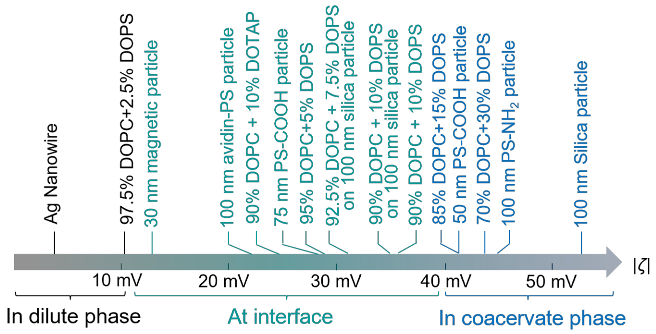


**Figure S4. Relationship between partitioning of particles across coacervates with absolute particle zeta potential.**

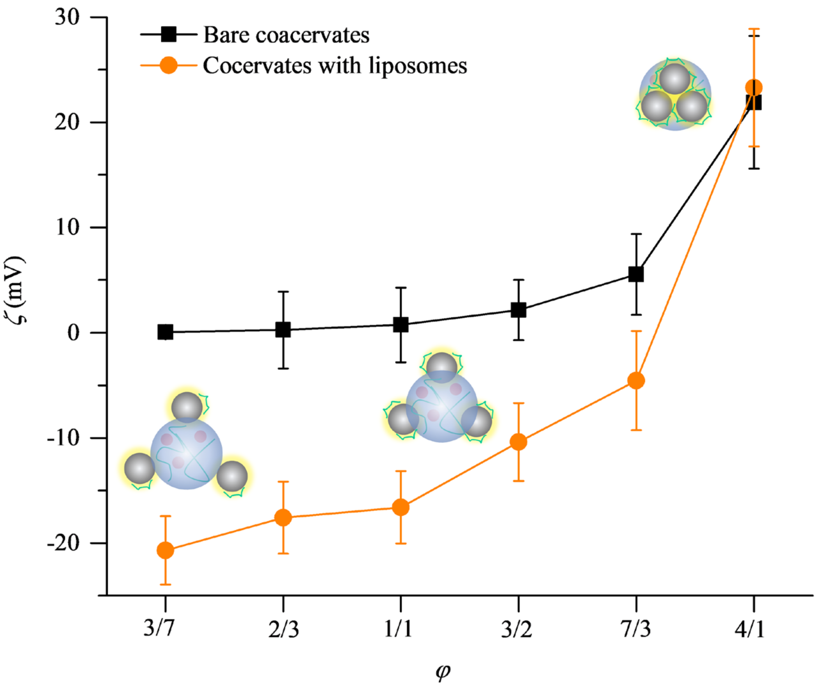


**Figure S5. Zeta potential $\zeta$ of bare coacervates and coacervate-liposome hybrid structures with $\varphi$ in PDDA-ATP- DOPC/DOPS ($P$=10%) liposome system.** The schematic is the structure of coacervate-liposome hybrid structure depending on $\varphi$. More balanced adsorption of PDDA on liposome is indicated with increasing $\varphi$ from the $\zeta$ of coacervate-liposome structures. This facilitates larger $\gamma_{p\text{-}d}$ and consequently partitioning preference of liposome for coacervate phase as observed in experiments.

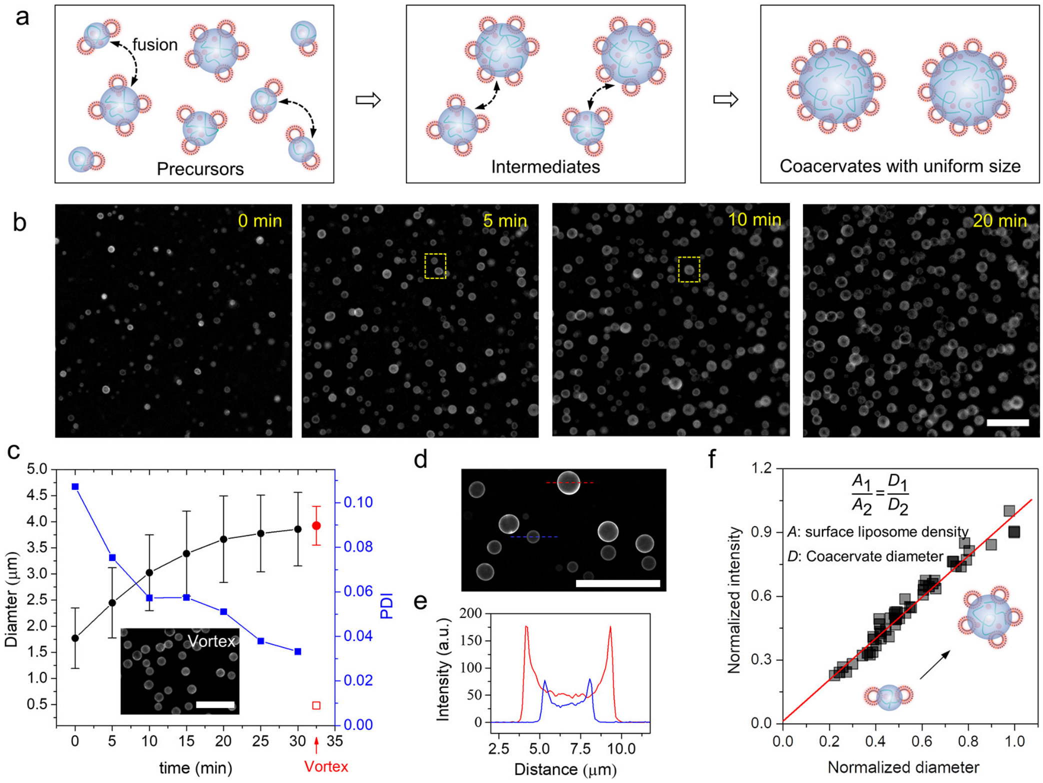


**Figure S6. Formation of liposome stabilized PDDA-ATP coacervates through free coalescence. a**, the schematic illustrating the formation of liposome stabilized coacervates via the coalescence of small coacervates. **b**, fluorescence images of the sample at 0 min, 5 min, 10 min and 20 min. The coacervate in the dash box at 10 min is formed via fusion of the two coacervates in the dash box at 5 min. **c**, variation of coacervate diameter (black line) and PDI (blue line) via free coalescence. For comparison, the diameter (red solid circle) and PDI (red hollow box) of coacervates formed via vortex are provided. The inset is the fluorescence image of coacervates formed via vortex. **d**, fluorescence image of the sample during the fusion process containing coacervates with different size. **e**, the profile of the fluorescence intensity along the dash line in **d**. Larger coacervates displayed higher fluorescence intensity, indicating denser packing of liposomes during the coalescence process. **f**, statistical analysis of the fluorescence intensity and diameter of coacervates from 50 coacervates in **d**. A linear increase of fluorescence intensity with coacervate diameter is observed indicating the increased liposome packing density on coacervates during the fusion process. The scale bars are 20 μm. The error bar represents the standard deviation. For all cases, $n \geq 100$.

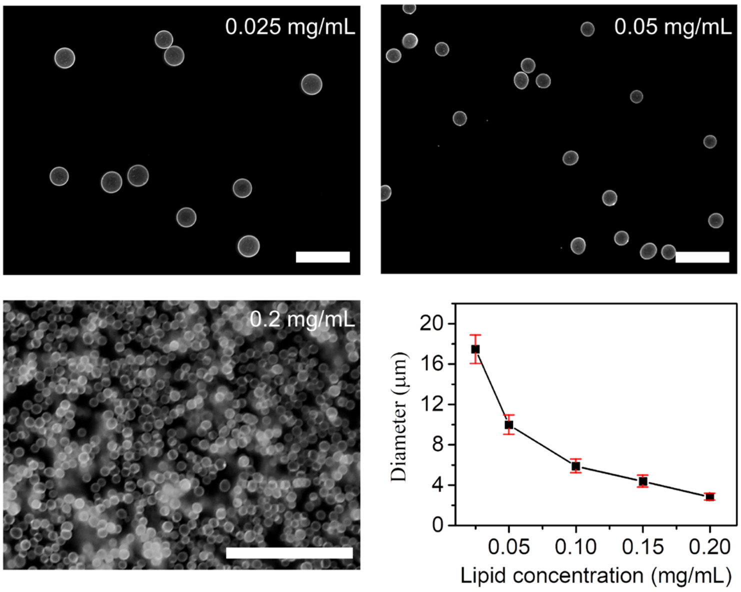


**Figure S7. Influence of 90% DOPC + 10% DOPS liposome concentration on coacervate diameter ($\varphi$=13/7).** The scale bars are 50 μm. Coacervate diameter decreases with the increased liposome amount. The error bar represents the standard deviation. For all cases, $n \geq 100$.

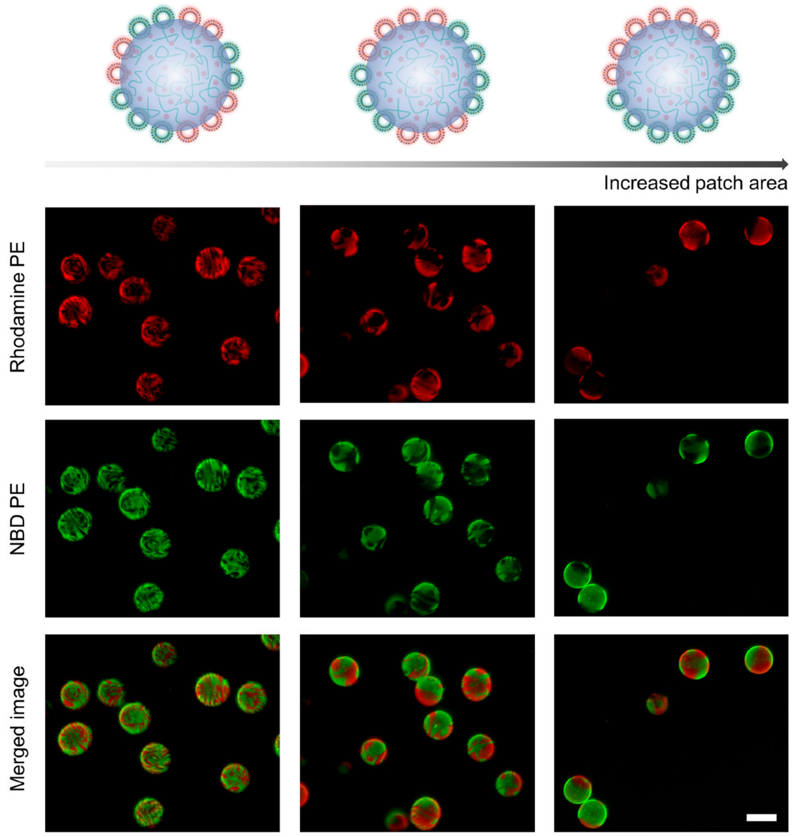


**Figure S8. Patch size control of PDDA-ATP ($\varphi$=3/2) coacervates with heterogeneously distributed G-liposome and R-liposome at surface.** The scale bar is 10 μm.

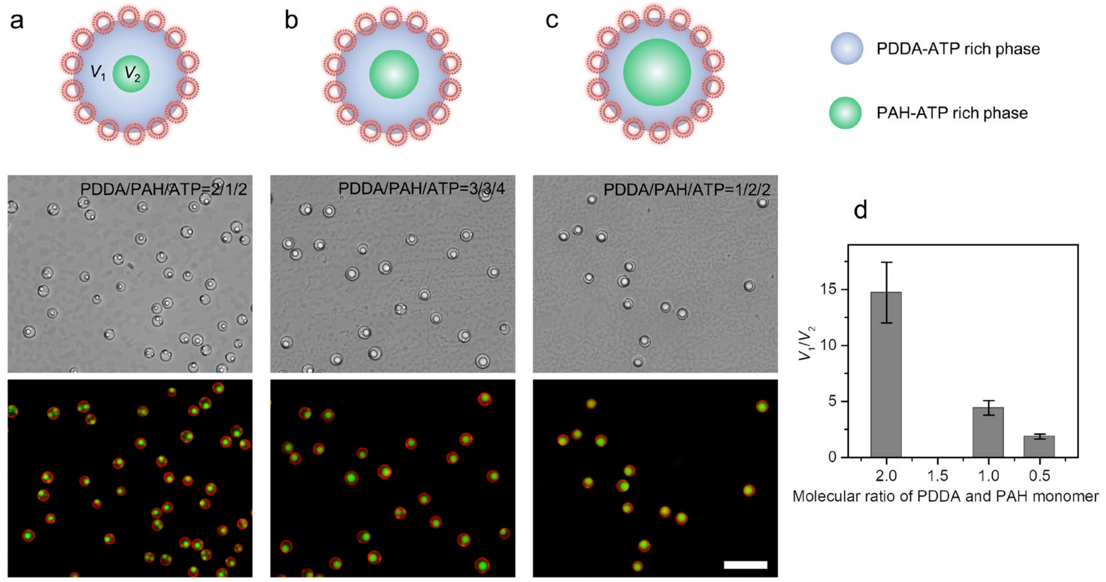


**Figure S9. Formation of liposome stabilized multiphase coacervates with various volume ratios ($V_1/V_2$) of PDDA-ATP rich phase ($V_1$) to PAH-ATP rich phase ($V_2$). a-c**, the schematics, bright field images and fluorescence images for liposome stabilized coacervates obtained at PDDA/PAH/ATP ratios of 2/1/2 (**a**), 3/3/4 (**b**), and 1/2/2 (**c**). **d**, relationship of the calculated $V_1/V_2$ with PDDA/PAH molecular ratios. The scale bars are 20 μm. The error bar represents the standard deviation. For all cases, $n \geq 100$.

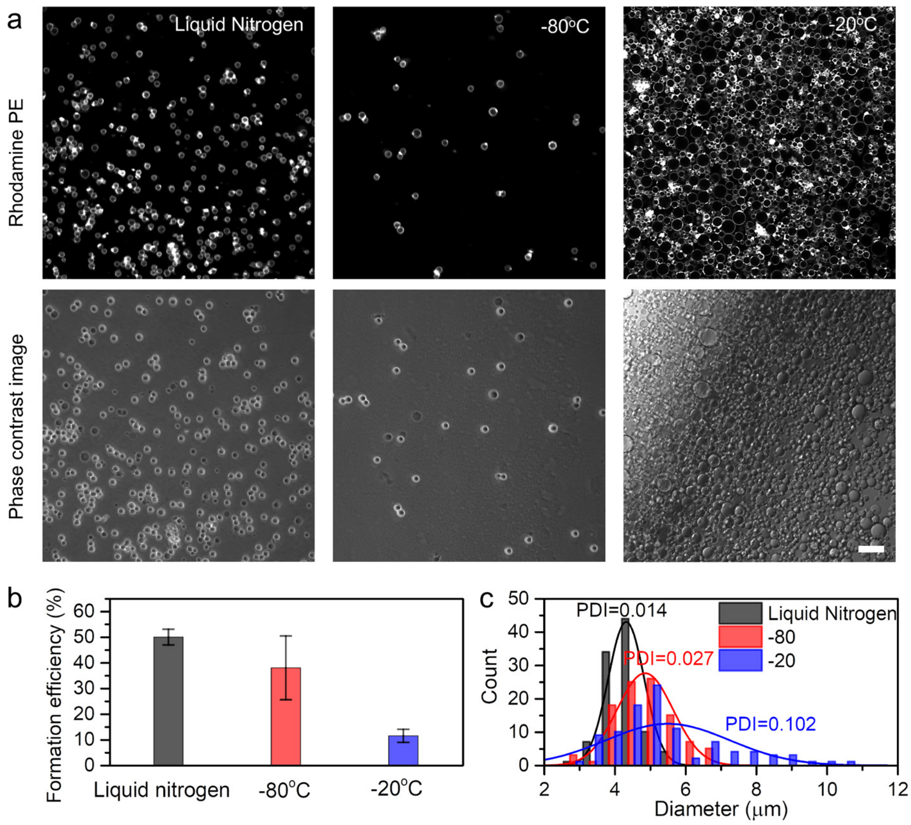


**Figure S10. Coacervate-GUVs formation at different freeze condition.** **a**, Fluorescence and phase contrast images of coacervate-GUVs formed via freezing in liquid nitrogen, at -80ºC, and at -20ºC. Lipid concentration is 0.025 mg/mL. **b**, Formation efficiency of coacervate-supported GUVs from coacervates coated by liposome particles at different freezing temperature. **c**, Distribution of coacervate-GUVs formed at different freeze condition. The scale bar is 20 μm. The error bar represents the standard deviation. For all cases, $n \geq 100$.

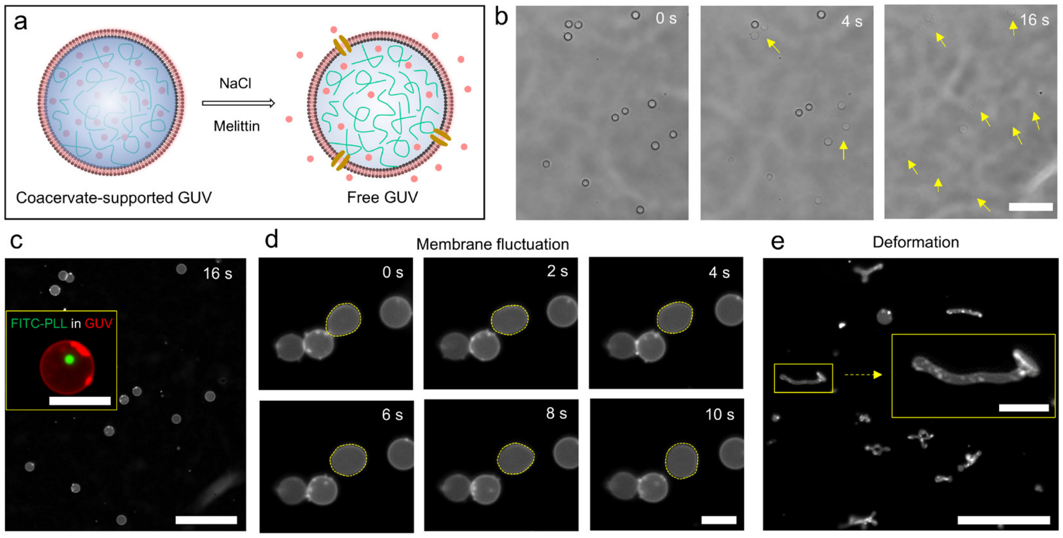


**Figure S11. Coacervate nucleus dissociation in coacervate-supported GUVs.** **a**, Schematic showing the transformation of coacervate-supported GUV into free GUV at the existence of 150 mM NaCl and 0.5 μg/mL melittin. **b**, Bright field image showing the dissociation of coacervate. **c**, Fluorescence images of free GUVs. The inset is a merged image of GUV (red fluorescence) with FITC-PLL-ATP coacervate (green fluorescence) inside when FITC-

PLL isencapsulated. Although PDDA-ATP coacervate dissociated at the existence of 150 mM NaCl and 0.5 μg/mL melittin, FITC-PLL forms a small coacervate with ATP. **d**, Fluorescence images showing membrane fluctuation of free GUVs. The yellow dash line illustrates the outline of a GUV. **e**, Fluorescence images of the formed lipid tubes from free GUVs after the addition of 2 M glucose. The scale bars in **b**, **c** and **e** are 50 μm. The scale bars in **d**, the inset in **c** and **e** are 50 μm.

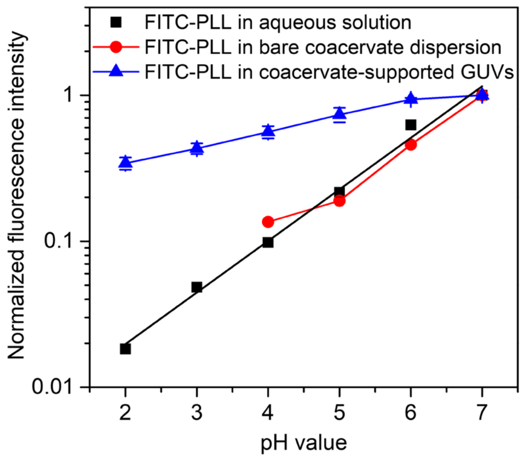


**Figure S12. Relationship of the fluorescence intensity *I* of FITC-PLL in bulk solution, coacervate, and CGUV with the pH value of dispersion media**. A linear relationship between log (*I*) and pH value of dispersion media is observed for FITC-PLL in bulk solution and coacervate. Based on it, we estimate the pH value in CGUV.

1 Olson, F., Hunt, C. A., Szoka, F. C., Vail, W. J. & Papahadjopoulos, D. Preparation of liposomes of defined size distribution by extrusion through polycarbonate membranes. *Biochimica et Biophysica Acta (BBA) - Biomembranes* **557**, 9-23, doi:https://doi.org/10.1016/0005-2736(79)90085-3 (1979).
